# Engineering Selective Molecular Tethers to Enhance Suboptimal Drug Properties

**DOI:** 10.1101/2020.02.01.929356

**Authors:** Alan B. Dogan, Horst A. von Recum

## Abstract

Small-molecule drugs are utilized in a wide variety of clinical applications, however, many of these drugs suffer from one or more suboptimal property that can hinder its delivery or cellular action in vivo, or even shelf an otherwise biologically tolerable drug. While high-throughput screening provides a method to discover drugs with altered chemical properties, directly engineering small-molecule bioconjugates provides an opportunity to specifically modulate drug properties rather than sifting through large drug libraries with seemingly ‘random’ drug properties. Herein, we propose that selectively “tethering” a drug molecule to an additional group with favorable properties will improve the drug conjugate’s overall properties, such as solubility. Specifically, we outlined the site-specific chemical conjugation of rapamycin (RAP) to an additional “high-affinity” group to increase the overall affinity the drug has for cyclodextrin-based polymers (pCD). By doing so, we found that RAP’s affinity for pCD and RAP’s window of delivery from pCD microparticles was tripled without sacrificing RAP’s cellular action. This synthesis method was applied to the concept of “affinity” for pCD, but other prosthetic groups can be used in a similar manner to modify other drug properties. This study displays potential for increasing drug delivery windows of small-molecule drugs in pCD systems for chronic drug therapies and introduces the idea of altering drug properties to tune polymer-drug interactions.

## INTRODUCTION

While antibody and protein therapeutics have made significant advances in the past decade, small-molecule drugs remain a major player in today’s clinically-relevant targeted therapeutic treatments, representing a multi-billion-dollar market. Directly acting upon intracellular molecular pathways, small-molecule drugs have been widely used in applications ranging from wound healing to combination cancer therapies (e.g. apixaban, lenalidomide, rivaroxaban). However, discovery of novel small-molecule drugs is greatly limited by drug discovery methods, many of which rely on natural sources, take years to isolate and develop and produce molecules with fixed physical and chemical properties. While high-throughput molecular screening methods help alleviate this time-strain, screening large libraries of drug analogs is many times a “shot in the dark” [1].

Small-drug bioconjugation and engineering shows promise to enhance suboptimal drug properties premeditatedly, making good drugs into great drugs, and bypassing the need for robust molecular libraries. These strategies have been applied to large-molecule drugs (e.g. proteins) for decades. For many smallmolecule drugs, structure and function are directly correlated, and non-specific chemical alteration can attenuate or eliminate cellular action. To prevent this loss of function, biocatalysts can be utilized in order to favorably target specific functional groups outside the protein-binding domains of the drug [2].

Sirolimus (Rapamycin; RAP), a commonly-used small-molecule mTOR inhibitor, is notoriously “tricky” to conjugate and modify, as much of its chemical structure is involved in binding to its target, FK506 binding protein, which impacts downstream processes involving cell survival, migration, and cell-cell signaling [3]. This leaves the sterically-hindered hydroxyl group on C40 as the optimal location for modification, and many have taken advantage of this by synthesizing “rapalogs” (Rapamycin analogs) including Temsirolimus and Everolimus [4]. Mainly, these modifications induce limited changes in solubility and in vivo half-life and rapalogs still require daily oral or IV injection for sufficient systemic treatment [4]. Even worse, for immunosuppression and cancer treatments, bolus, systemic administration of RAP and rapalogs can result in adverse events including lowered platelet counts [5], pulmonary complications [6], and cardiac irregularities [7].

Efforts have been made to increasingly localize RAP therapies by utilizing polymer-mediated delivery systems, however these attempts are largely limited by the maximum loading of drug into the polymer matrices and release profiles of usually much less than 20 days [8]. As many patients require treatment on the scale of weeks to months, there is a need to increase the time duration of polymer-controlled release of RAP.

Our group has previously shown that the loading and release of RAP can be improved utilizing polymerized cyclodextrins (pCD) [9]. pCD hydrogels have previously shown to successfully deliver small-molecule drugs on the scale of a few weeks leveraging thermodynamic interactions, or ‘affinity’, between hydrophobic molecules and cyclodextrin’s hydrophobic complexation domain, or “pocket” [10,11,12,13,14]. pCD networks can be made into large macrostructures including disks and particles, which have been shown to load relatively high amounts of drugs while altering the window of release, resulting in an extended ‘affinity-based’ release rather than a purely diffusion-based release [15,16]. This system produced a 28-day window of release, which satisfies the time frame for wound healing and cancer therapeutic delivery [17] but not for increasingly chronic applications like immunosuppression [9].

We propose to improve RAP’s suboptimal affinity towards cyclodextrin complexation by selectively tethering the drug to either a “high-affinity” group, adamantane, or an additional RAP molecule to enhance its affinity for pCD hydrogels. Traditionally, small-molecule drugs interact with CD pockets in a 1:1 fashion – one ligand is included in one CD “host”. Increasingly complicated interactions, such as a 1:2 ligand:host interactions and polymer “threading” through CD pockets, can be leveraged to increase these “affinity” interactions [18,19]

Through the addition of an additional “affinity” group, Adamantane has previously been used as a high-affinity “anchor” for other drugs like doxorubicin, however region-specific conjugation was not achieved [20,21]. Herein, we outline a broadrange synthesis for covalently tethering small-molecule drugs to additional groups to enhance suboptimal properties. Explicitly, we showed that if RAP, used as a model drug, is region-specifically tethered to adamantane or an additional RAP molecule with a poly(ethylene) glycol (PEG) spacer, then the conjugate will have enhanced affinity towards pCD hydrogels, ultimately leading to enhanced loading and extended release, all without loss of biofunctionality. The dimeric version of RAP also used the PEG spacer allowing each RAP domain to separately interact to a CD group. As shown in **Figure 1,** we hope to leverage a 1:2 ligand:host interaction to increase the thermodynamic interactions between RAP and pCD. This is different from a dimeric analogue version of FK506, known as FK1012, which lacks a flexible hinge region, reducing the capacity to interact with two CD groups [22].

**Figure 1:**
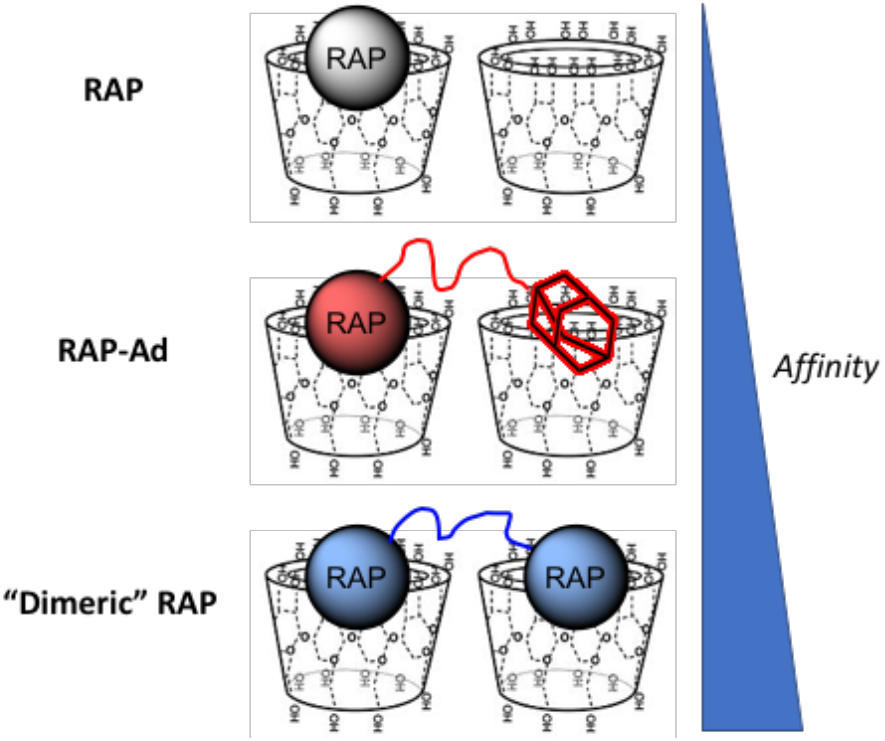
RAP traditionally complexes with cyclodextrin 1:1 (ligand:host). The conjugation of additional “affinity” groups to RAP is expected to improve affinity by leveraging multiple complexations in pCD hydrogels, as both 1:1 and 1:2 interactions will occur.

## MATERIALS AND METHODS

β-cyclodextrin (β-CD) prepolymer, lightly crosslinked with epichlorohydrin, was purchased from Cyclolab (Budapest, Hungary). HyperSep C18 SPE Cartridges was purchased from Sigma-Aldrich (St. Louis, MO). Sirolimus (RAP) was purchased from Biotang (Lexington, MA). Propargyl-PEG1000-amine and PEG3000-diamine was purchased from BroadPharm (San Diego, CA). Cuprisorb was purchased from Seachem Laboratories (Madison, GA). 7” NMR tubes were purchased from Bruker (Billerica, MA). Phenomenex Luna 5u C18 250×4.6 mm column was purchased from Phenomenex (Torrance, CA). All other reagents, solvents, and chemicals, including dialysis bags were purchased from Fisher Scientific (Hampton, NH) in the highest grade available.

**Figure 2:**
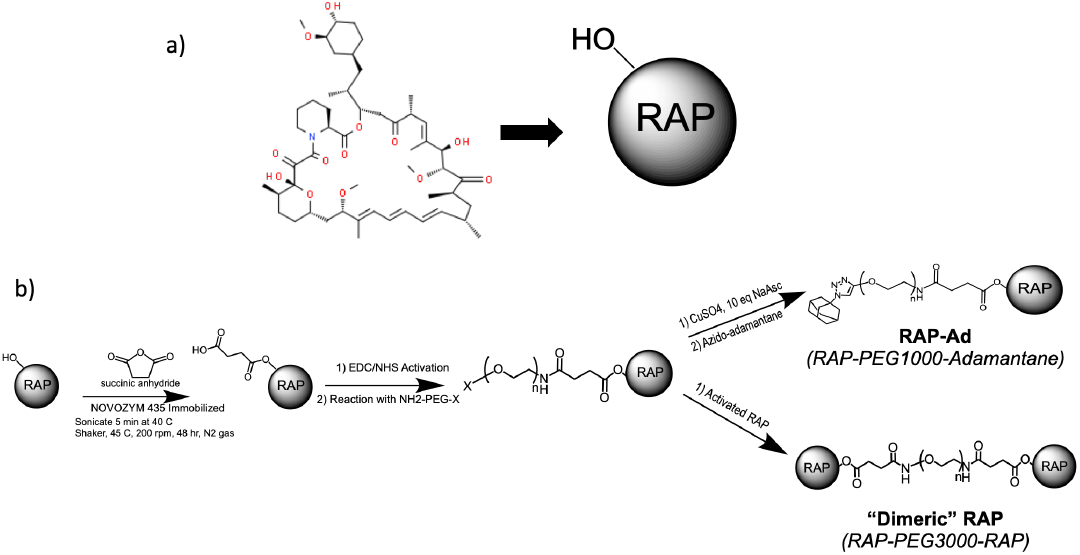
a) Molecular structure of Rapamycin with cartoon representation isolating hydroxyl group on C40. b) Synthesis outline for RAP-Ad and “Dimeric” RAP. X functional group represents an lkyne for Cu(II) ‘click’ chemistry and an amine for EDC/NHS coupling.

Small-molecule drug modifications were modeled using RAP conjugation with a PEG spacer. Briefly, the synthesis of RAP-Ad (RAP-PEG1000-Adamantane) and “Dimeric” RAP (RAP-PEG3000-RAP) follows lipase B mediated RAP succinylation, EDC/NHS coupling of RAP to bifunctionlized PEG-amine, and azide-alkyne Huisgen cycloaddition (CuAAC ‘click’ chemistry) of azido-adamantane to PEG’s terminal alkyne or additional conjugation of EDC/NHS activated RAP conjugation, respectively.

### Selective RAP Succinylation

Due to RAP’s functionality as a mTOR (mammalian target of rapamycin) protein inhibitor, it has been found that only specific modification of C40 yields RAP analogs with retained biofunctionality. Therefore, immobilized Candida antarctica lipase B (NOVOZYM 435 Immobilized) is used to hold RAP in a conformation in which only the C40 hydroxide group is exposed [23]. Briefly, 250 mg of NOVOZYM 435 Immobilized was added to a 1.85:1 mass ratio of RAP to succinic anhydride in 4:1 toluene:dichloromethane and allowed to react for 48 hours at 37 C in an inert, N_2_ atmosphere at 200 rpm. The solvent was degassed for 5 minutes before the reaction took place and subsequently sonicated for 5 minutes at 40 C to ensure solubilization of reactants.

After completion, the enzyme was filtered off with dichloromethane. The solution was then filtered through a silicon column with dichloromethane:methanol (13:1) as eluent to remove residual succinic anhydride.

### EDC/NHS Coupling

Following the succinylation of RAP, the free carboxylic acid group was activated through EDC/NHS chemistry. Breifly, 10 molar excess EDC (1-Ethyl-3-(3-dimethylaminopropyl) -car-bodiimide) and 4 molar excess NHS (N-hydroxysuccinimide) was reacted with succinylated RAP in DMSO:NaHCO_3_ (pH 8.3, 1:1.4 v/v) with excess EDC/NHS for 5 hr at room temperature while stirred. Once reacted, excess EDC/NHS was removed by running the solution through a HyperSep C18 Column. First the cartridge was washed with 10 mL methanol/water (90:10) and then 10 mL water as preconditioning, and 20 mg sample in 2 mL acetonitrile/water was loaded on column. Extraction was carried out by acetonitrile/water (70:30) as eluent. Several fractions (4 fractions, each 3 mL) were collected, and UV absorbance was checked at 280 nm. The fractions containing “activated” RAP was dried by lyophilization and used for further steps.

For RAP-Ad synthesis, EDC/NHS activated RAP was then reacted with 2 molar equivalents of Propargyl-PEG1000-amine at room temperature overnight in in DMSO:NaHCO_3_ (pH 8.3, 1:1.4 v/v). For “Dimeric” RAP synthesis, 0.5 molar equivalent of PEG3000-diamine was reacted under similar conditions.

### CuAAC ‘Clicked’ Adamantane

To functionalize the opposite end the PEG during RAP-Ad synthesis, an adamantane group was “clicked” to PEG’s free alkyne end using Copper-Catalyzed Azide-Alkyne Cycloaddition (CuAAC) chemistry. RAP-PEG1000-propargyl was reacted with 5 molar equivalents of azido-adamantane in the presence of 5 molar excess copper sulfate and 10 molar excess NaAsc ((+)-Sodium L-ascorbate) for 12 hours in DMSO:H_2_O (2:1). A green color was observed if sufficient Cu(II) ions was present during reaction.

To remove excess copper ions and other unreacted reagents after reaction completion, the sample dialyzed by a 1 kD cutoff molecular weight dialysis membrane for 48 hours in 1 Liter of deionized H_2_O (diH_2_O) solution with a 5 g bed of Cuprisorb to act as a sink for excess copper ions.

### “Dimeric” RAP Synthesis

To functionalize the second amine group of the RAP-PEG3000-amine conjugate, RAP-PEG3000-amine was reacted with two molar equivalents of EDC/NHS “activated” RAP overnight in in DMSO:NaHCO_3_ (pH 8.3, 1:1.4 v/v) with excess EDC/NHS. Reaction was then dialyzed by a 3.5 kD cutoff molecular weight dialysis membrane for 48 hours in 1 L of diH_2_O.

### Nuclear Magnetic Resonance

Nuclear magnetic resonance (NMR) was used to verify successful conjugation during synthesis steps. All spectra of presented chemical species was recorded by Bruker 300 MHz NMR system (Bruker, Germany) in DMSO-D_6_ solvent.

### Fourier Transform Infrared Spectroscopy

Fourier Transform Infrared Spectroscopy (FT-IR) was used to verify bond formation during RAP succinylation. FT-IR spectrums were obtained with a Bio Rad Digilab FTS 3000-MX Excalibur Series FT-IR. Briefly, samples were mixed with potassium bromide (KBr) in a 30:70 w/w ratio. 13 mm pellets were formed by compacting the powder under 6-7 metric tons for thirty seconds.

### pCD Microparticle Synthesis

pCD polymers were synthesized into microparticles as example drug delivery vehicles. To make the microparticles, epichlorohydrin-crosslinked β-cyclodextrin prepolymer was solubilized in 0.2 M potassium hydroxide (25% w/v) and heated to 60 C for 10 minutes. Light mineral oil was warmed in a beaker with a Tween85/Span85 solution (24%/76%) and stirred at 500 RPM. Ethylene glycol diglycidyl ether was added drop-wise and the solution was vortexed for 2 minutes before pouring into the beaker with the oil/Span/Tween85 mixture, increasing temperature to 70°C, and stir speed for 3 hours. The microparticles were then centrifuged at 200 x g to be separated from the oil mixture, washed with excess hexanes twice, excess acetone twice, and finally de-ionized water (diH2O) twice. The microparticles were frozen and lyophilized before further use.

### Drug Loading/Release

Drug loading and drug release kinetics of RAP, RAP-Ad, and “Dimeric” RAP in pCD microparticles were tested in vitro. Briefly, 100 ug of pCD was incubated in concentrated drug solutions (either ‘low’ [3.7 mM] or ‘high’ [21 mM] conditions) in DMSO for 72 hours. Particles were spun down (10,000 RPM) and incubated in 500 uL of “physiological release buffer” (phosphate buffered saline [PBS], 0.1% Tween80). At recorded time points, the release buffer was removed and replaced with fresh buffer to better mimic in *vivo* conditions: each sample’s absorbance was measured at 278 nm to determine drug concentration based on standard curves.

### Affinity Testing

Favorable drug loading and release kinetics in pCD polymers are assumed to be leveraged from affinity interactions between host and ligand. To confirm that “affinity” is responsible for polymer-drug interactions, we tested the thermodynamic interactions between pCD and RAP/RAP-conjugates in simulation and experimentally.

#### Docking Simulation

To test “affinity” we utilized a computer “docking” software to predict thermodynamic interactions between host and ligand based on spatial data. Molecular structure data files for RAP and β-CD were downloaded from the PubChem database. Structures were converted to PDBQT format and energy was minimized before loading RAP as a ligand and cyclodextrins as a host in PyRx (Molecular Graphics Laboratory, The Scripps Research Institute, La Jolla, CA). The Autodock Vina algorithm was used to predict the strength of the ligand/guest interaction.

#### Surface Plasmon Resonance

To obtain “affinity” experimentally between β-CD monomers and RAP, RAP-Ad, and “Dimeric” RAP was measured experimentally through surface plasmon resonance (SPR) with a Biacore X100 system (GE Healthcare Bio-Sciences, Pittsburgh, PA) according to previous protocols [9,21]. The surface of a sensor chip CM-5 was conjugated with EDC (0.4 M) and NHS (0.1 M) followed by 10 mM 6-amino-6-deoxy-β-cyclodextrin (CycloLab) suspended in HBS-N buffer (a HEPES balanced salt solution with pH 7.4). The other channel was conjugated similarly with amino-dextran (Thermo Fisher Scientific) to determine specific versus nonspecific interactions with a chemically similar but non-affinity substrate. The remaining functional groups were capped with ethanolamine. A multi-cycle kinetic experiment was performed with drug dissolved in a 1% dimethyl sulfoxide MilliQ water solution and was regenerated with 100 mM sodium hydroxide between samples. The differential responses between the channels were fit to both steady state affinity using Biacore evaluation software. Reported K_D_ values were all within model confidence interval (Chi^2^ values below 10% of the maximum SPR response) [24].

### Cell Migration “Scratch” Assay

As the mTOR pathway is responsible for regulating cellular migration, a fibroblast cellular migration assay was used as an indirect indicator for mTOR inhibition. PT-K75 porcine mucosal fibroblasts (ATCC, Manassas, VA) were cultured in Dulbecco’s Modified Eagle’s Medium (DMEM) supplemented with 15% fetal bovine serum (FBS) and 1% penicillin/streptomycin (P/S) at 37°C, 5% CO_2_. Cells were plated at ~19,200 cells per well and allowed to adhere for 2 hours in standard media, after which standard media was replaced with “starvation” media (DMEM, 0.5% FBS, 1% P/S) and incubated for 24 hours. Plates were then “scratched” with a sterile 200 uL pipette tip, washed with “starvation” media, imaged, and treated with drug aliquots for 25 hours. At 25 hours, scratches were reimaged and compared to t0 and normalized to buffer-only controls. Images were analyzed in ImageJ with a variance-based macro to measure total area of “wound”.

### Cell Proliferation Assay

mTOR signaling pathway also plays a vital role in cell metabolism and proliferation. Therefore, cellular proliferation is closely linked to mTOR activation. PT-K75 porcine mucosal fibroblasts were cultured according to previously stated methods. Cells were plated at around 7,000 cells per well and allowed to adhere in growth media for 2 hours. Aliquoted concentrations of each drug was added to wells (n=3) and incubated for 24 hours. According to manufacturer protocols, 10 μl of 0.15 mg/ml resazurin, which is used to quantify fibroblast metabolic activity, viability and proliferation [9,22], was added per treatment well. After incubation for an additional 21 hours, fluorescence (530/590 excitation/emission) was measured with a Syngergy H1 Microplate Reader (BioTek Instruments, Inc., Winooski, VT) and normalized to buffer controls.

### Statistical Analysis

Statistics for drug loading, drug release, cellular migration and cellular proliferation were calculated in Origin (OriginLab, Northampton, MA). Statistically significance was defined as p < 0.05 with further specifications stated in figure captions.

## RESULTS AND DISCUSSIONS

### Succinylation of RAP

To allow selective conjugation of PEG to RAP, RAP succinylation must first be confirmed to introduce a carboxylic group for EDC/NHS activation. Successful RAP succinylation was confirmed by FT-IR and 1H-NMR spectroscopy. The addition of an observable peak at 1740 cm^-1^ and a broadened OH peak from 2750 to 3100 cm^-1^ suggests the hydroxyl group on the unmodified Sirolimus has been replaced with a carboxylic group (-COOH), matching the description of succinylated RAP outlined in Behrouz H *et* al [23] (**Figure 3a**).

**Figure 3:**
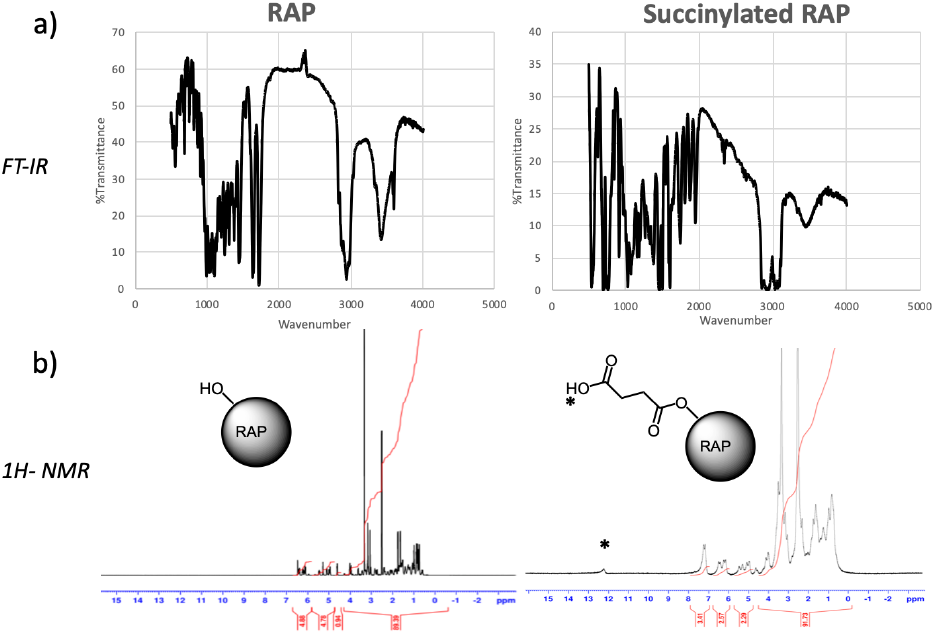
**a)** FT-IR of RAP and Succinylated RAP. v = 1740 (C=O), 3100 (broadened O-H) cm^-1^ suggests terminal carboxylic acid formation. **b)** 1H NMR (DMSO-d_6_). [*] δ = 12.1 ppm (-COOH) corresponding to addition of acidic hydrogen after succinic anhydride opening.

Succinylated RAP was further observed 1H NMR – the introduction of δ =12.1 ppm corresponds to the acidic hydrogen found after the opening of succinic anhydride during the reaction, suggesting succinylation was successful at a ratio of about 1:1 (-COOH:RAP) according to chemical shift peak integration (**Figure 3b**).

Overall succinylation efficacy range was 73-85%, similar to values previously reported, calculated equation 1 (molecular weight for RAP and succinylated RAP assumed to be 914.172 g/mol and 1000 g/mol, respectively) [23]

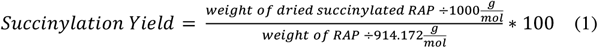

### EDC/NHS Coupling to PEG

After the introduction of a carboxylic group to RAP, we aimed to conjugate succinylated RAP to the free amine group on bifunctional PEGs. The intermediate EDC/NHS coupling of PEG to RAP was also confirmed with 1H NMR. EDC/NHS coupling success can be seen in the disappearance of δ =12.1 ppm peak (COOH) and the introduction of a peak at δ =7.8 ppm, corresponding to the amine peak (NH2) of EDC (**Figure 4**). Successful conjugation of PEG1000 to RAP can be seen from the disappearance of the δ =7.8 ppm EDC peak and the introduction of a large peak at δ =3.6 ppm corresponding to the (CH2-CH2-O) repeat units of PEG. The disappearance of the 8 ppm peak after conjugation suggests RAP coupling progressed and the remaining species after dialysis contained both RAP and PEG characteristics.

**Figure 4:**
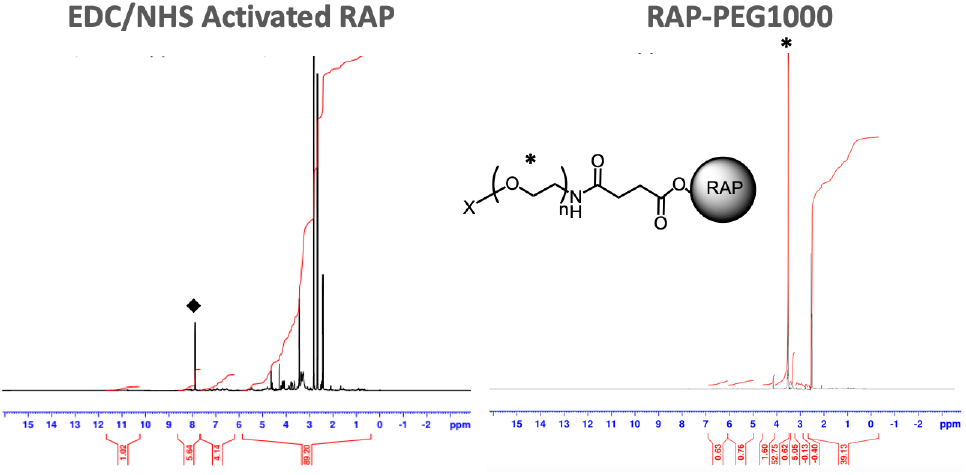
1H NMR (DMSO-d_6_) EDC/NHS coupling: δ = 12.1 ppm (-COOH) peak replaced by [◆] δ = 8 ppm (amine bond). PEG1000 [*] δ = 3.6 ppm (CH2-CH2-O) repeat units of PEG.

### CuAAC ‘Clicked’ Adamantane

For RAP-Ad synthesis, we aimed to “click” an adamantane functional group to the free alkyne end of RAP-PEG1000-alkyne. Adamantane conjugation to the PEG1000 chain yielded two distinct peaks at δ = 1.6 & δ =1.7 ppm, corresponding to adamantane’s CH2 groups. In addition, the formation of the triazole ring resulted in a characteristic peak at δ = 7 ppm (**Figure 5**). Comparing the area of the PEG1000 and adamantane peaks, there appears to be approximately one adamantane for each PEG molecule. Adamantane conjugation was further confirmed via 13C NMR, with introduced peaks at δ = 29.35 and δ = 39.5 (*Supp. Fig. 1*).

**Figure 5:**
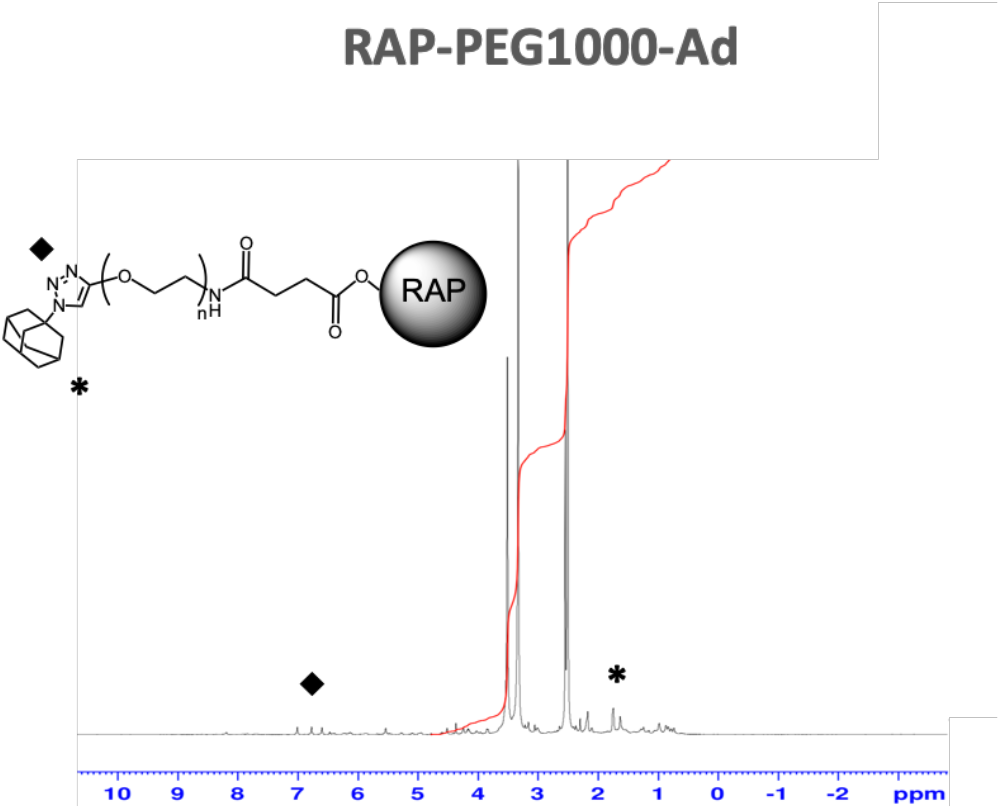
1H NMR (DMSO-d_6_) RAP-PEG1000-Ad: [*] dual adamantane δ = 1.7 & 1.6 ppm, [◆] tetrazole ring peaks δ ~ 7 ppm.

### “Dimeric” RAP Synthesis

To complete the synthesis of “Dimeric RAP”, we sought to conjugate an additional “activated” RAP molecule to the free amine end of RAP-PEG3000-amine. Conjugation of an additional RAP molecule to RAP-PEG3000-amine via EDC/NHS coupling was confirmed via 1H NMR and HPLC UV-Vis (Beckman HPLC, Phenomenex Luna 5u C18 250×4.6 mm column, λ= 280nm). EDC/NHS reaction efficiency heavily effected the conjugation yield and purification of the “Dimeric” RAP. As EDC/NHS coupling reactions have previously been shown to be improved by altering buffer type and pH, several alternative solvents were tested to see if the reaction could be pushed to favor increased conjugation [26]. The corresponding 1H NMR and HPLC UV-Vis of formulation that yielded the greatest conjugation of “dimer” (29.4% “Dimeric” RAP, 20.7% RAP-PEG3000) (**Figure 6**). Other attempts at increasing conjugation yield are show in *Supplemental Figure 2*.

**Figure 6:**
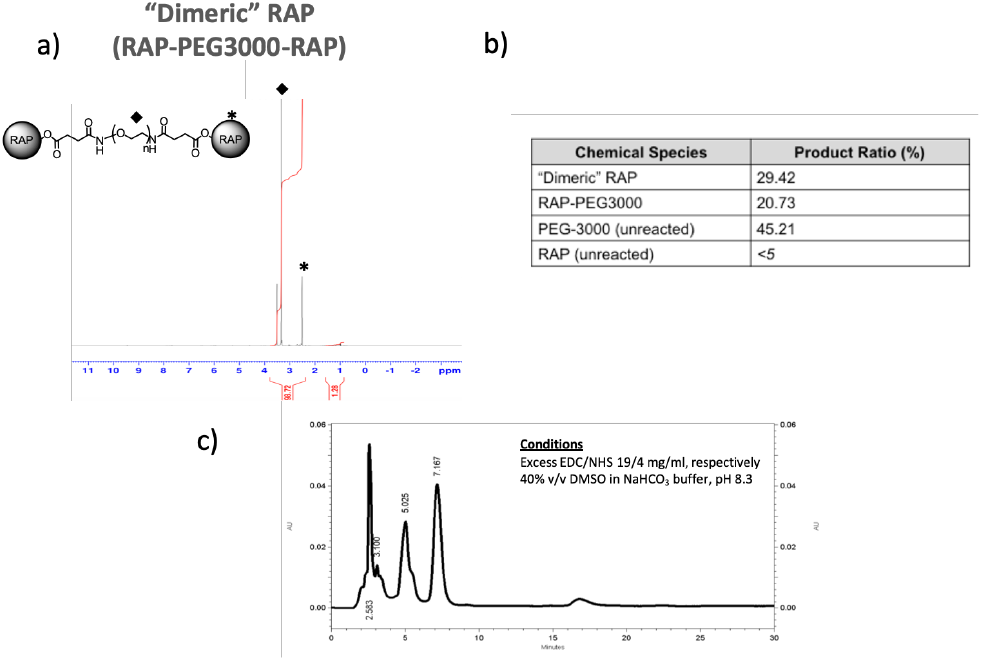
**a)** 1H NMR of “Dimeric” RAP: PEG3000 [◆] δ = 3.6 ppm peak and main RAP peak [*] δ = 2.6 ppm **b)** EDC/NHS reaction yield products determined from HPLC peak integral analysis **c)** Nonpolar-phase HPLC UV-Vis spectrum for “Dimeric” RAP synthesis.

### Molecular Tethers Increased RAP’s affinity for β-CD

Historically, cyclodextrin drug delivery leverages the “affinity” a drug (ligand) has for pCD’s inclusion complexes. Dissociation constants (K_D_) between small molecule drugs and cyclodextrins have previously been used to predict both loading efficiency and release profiles, as it is a representation of the thermodynamics of complexing drugs with cyclodextrin [12,18, 21]. RAP was previously reported to have an affinity of around 34 uM, which corresponded to a 28 day window of release [9]. A decrease in K_D_ suggests a higher affinity and subsequent increased loading and prolonged release. Both RAP-Ad (K_D_=12.41 uM) and “Dimeric” RAP (K_D_=11.82 uM) exhibited a significantly decreased K_D_ against β-CD monomers (**Table 1**).

**Table 1:**
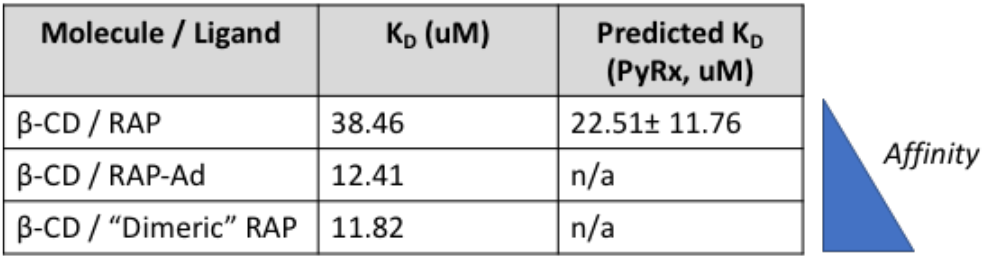
SPR kinetics results for unmodified and modified RAP against β-CD immobilized CM5 chip, 1% DMSO:diH2O running buffer. Reported K_D_ values were within the model confidence interval with Chi^2^ values below 10% of the maximum SPR response.

### Drug Loading and Drug Release Kinetics Were Both Enhanced in pCD Microparticles

In vitro drug loading and drug release studies have be used to predict future in vivo delivery efficacy. Two drug loading conditions were chosen: one “low concentration” in which the number of pCD complexation sites outnumbered the amount of drug, and one “high concentration” in which drug was added in large excess to pCD complexation sites. Here, we observed a significant increase in high concentration loading groups, and low concentration loading, with both RAP-Ad and “Dimeric” RAP loading nearly 17 and 6 times more efficiently than RAP, respectively (**Figure 7**).

**Figure 7:**
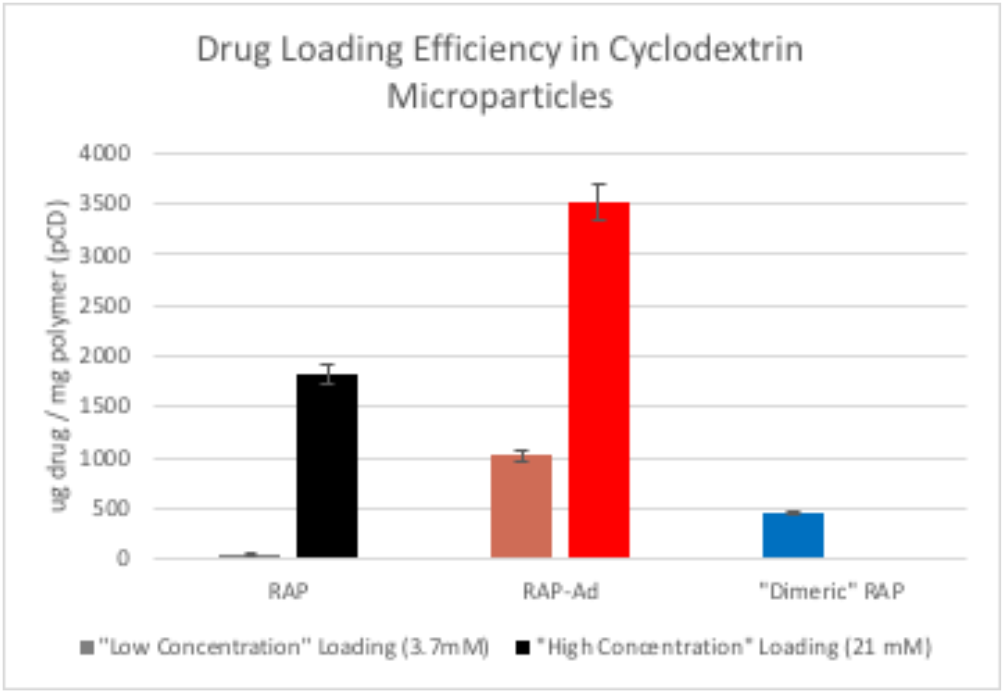
Total drug released after loading in β-CD microparticles with concentrations of 3.7 mM (“low concentration”) and 21mM (“high concentration”) [n=3]. Error bars represent the standard error of the mean.

All “low concentration” drug release profiles exhibited a limited “burst” release and released relatively consistently over time. RAP behaved similarly as previously reported, releasing over a timespan of around 20 days [9]. RAP-Ad and “Dimeric” RAP, however, both released to up to 65 days and had a reduced slope, correlating to the increase in affinity (**Figure 8a**). Additionally, the daily release of RAP-Ad was, on average, 6 times more concentrated than RAP control and “Dimeric” RAP was on average 2 times more concentrated than RAP control. This is a consequence of both increased affinity and drug solubility as a result of PEGylation (*Supp. Fig. 4*).

**Figure 8:**
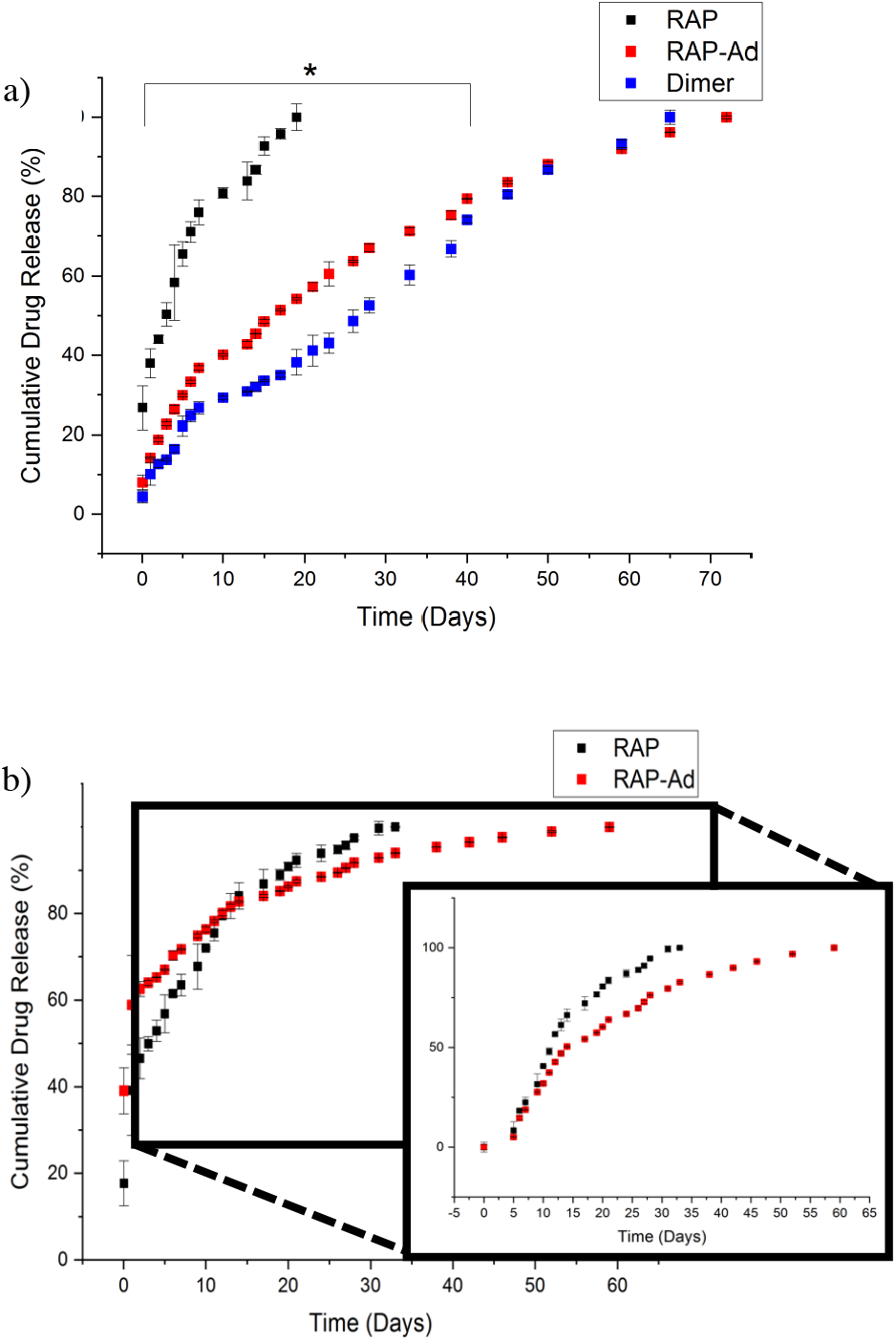
**a)** “Low concentration” (3.7 mM) cumulative release profile from β-CD microparticles for RAP, RAP-Ad, and “Dimeric” RAP. * All groups were statistically significant in first 40 days of release (p<0.05) by Student’s T-test. **b)** “High concentration” (3.7 mM) cumulative release profile from β-CD microparticles for RAP, RAP-Ad, and “Dimeric” RAP. Day 5-65 releases were renormalized to 100% release to highlight “affinity” segment of release.

“High concentration” releases for both RAP and RAP-Ad had a two-phase release: a solvent-exchange “burst” release within days 1-4 followed by an “affinity-based” release thereafter. Due to RAP-Ad’s higher affinity, drug delivery took place over a timespan of over an extended 60 days, similar to the release at low concentration while delivering comparable amounts of drug daily after day 20 (**Figure 8b**).

### Rapamycin Retains its Biofunctionality After Selective Modification

Previous attempts at creating modified RAP molecules (rapalogs) saw that non-specific modifications of RAP’s functional groups lead to loss of immunosuppressive properties [27]. Herein, we avoided this loss-of-function by utilizing a biocatalyst, lipase B, to direct conjugation to a site known not to interrupt protein binding. We utilized porcine fibroblasts as a model-mTOR pathway system, as fibroblasts have been shown to highly depend on mTOR for migration, proliferation, and survival [28,29]. Using a wound-healing “scratch” assay as a measure of mTOR activity (**Figure 8**) we found no significant differences in drug activity between RAP and RAP-Ad, while “Dimeric” RAP seemed to have diminished activity at lower concentrations (< 2 uM). We also found that RAP functioned comparably to both RAP-Ad and “Dimeric” RAP when inhibiting proliferation, with both groups showing improved proliferation prevention at concentrations lower than 1 uM (**Figure 9**).

**Figure 9:**
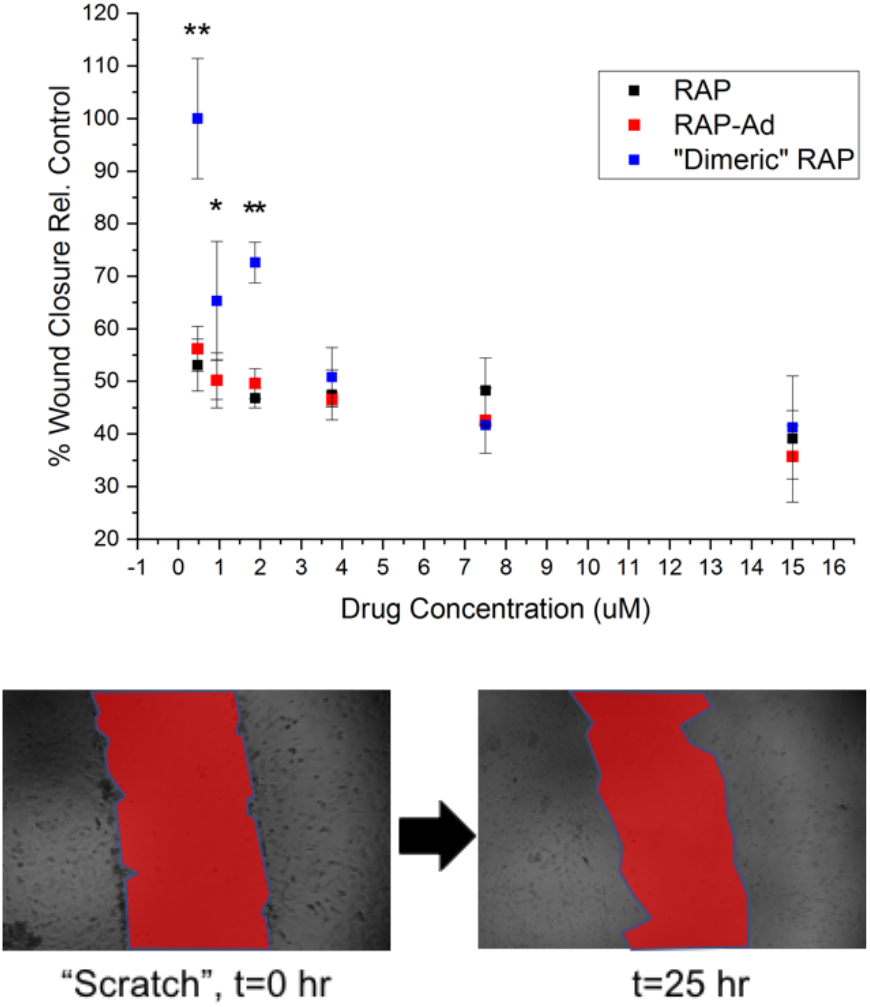
Scratch assay wound closure of PT-K75 porcine mucosal fibroblasts after 24 hour RAP, RAP-Ad, or “Dimeric” RAP treatment relative to PBS controls in serum starvation media. Area of each “scratch” was analyzed at time 0 and 25 hours utilizing ImageJ. Areas were compared to manual ROI selection for confirmation [n=3]. * indicates p<0.05 and ** indicates p<0.001 by twoway ANOVA with Tukey test.

### Discussions

This work demonstrates that RAP, a small-molecule drug, was specifically modified keeping its mTOR inhibitory activity in vitro (**Figures 9 and 10**) while greatly increasing its affinity for pCD hydrogels (**Table 1**). By tethering additional groups to RAP - either an adamantane or additional RAP molecule - we successfully synthesized site-specific bioconjugates that demonstrated enhanced loading and had an extended delivery of up to 72 days in pCD microparticles (**Figures 7 and 8**). More specifically, we saw a 3.6 fold window of delivery increase from many drug-delivery polymer systems and a 2.5 fold increase from RAP’s “affinity” release in pCD [9]. As the difference in K_D_ between RAP and RAP-Ad / “Dimeric” RAP was around 3-fold, this confirms a direct relationship between “affinity” and timespan of release [12].

**Figure 10:**
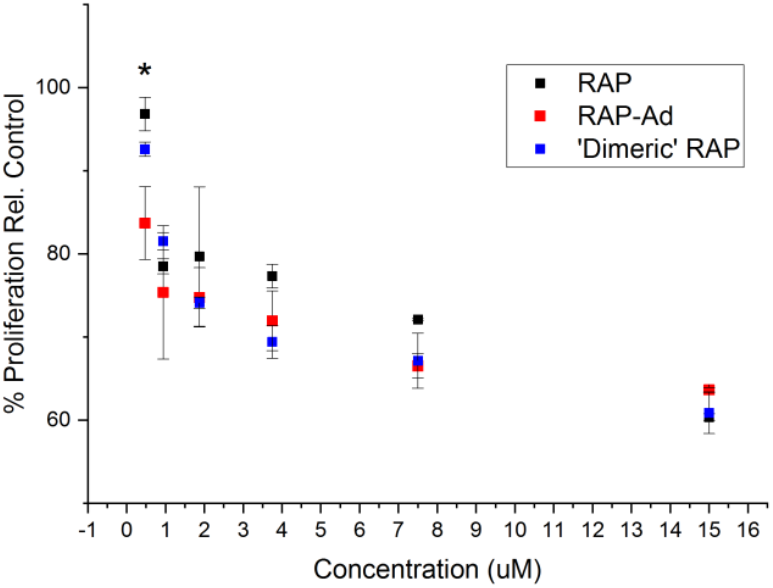
Proliferation of PT-K75 porcine mucosal fibroblasts after incubation with RAP, RAP-Ad or “Dimeric” RAP for 24 porcine mucosal fibroblasts hours. Fluorescence was compared between groups [n=3] and control after 28 hours of Alamar blue incubation. * indicates p<0.05 by two-way ANOVA with Tukey test.

PEG “threading” could also account for the prolonged release times for both RAP-Ad and “Dimeric” RAP. While PEG itself is predicted to have relatively low affinity for cyclodextrins, longer PEG chains have been shown to weave between pCD inclusion “pockets”, resulting in an increasingly crystalline structure [27,28]. It was observed that pCD microparticles loaded with either RAP-Ad and “dimeric” RAP precipitated quicker and were more opaque than RAP-loaded pCD, which is most likely a ramification of PEG-pCD threading. While this phenomena may not be contributing significantly thermodynamically, it may help explain the nonlinear release profile of “dimeric” RAP after day 10 (**Figure 8a**).

The “burst” release shown in the “high concentration” loading was predicted to be an artifact of excessive drug loading, where excess RAP molecules were not complexed with pCD but interspersed between pCD polymer chains. While this may seem disadvantageous for applications such as immunosuppression, other drugs, such as antibiotics require a “bolus” dose followed by continuous treatment [29]. With an understanding that the “burst” release from pCD hydrogels occur within 1-3 days, we can engineer sustained drug delivery systems with an inherent “burst” release response.

The performance of RAP-Ad and “Dimeric” RAP raises a few questions regarding the specific cellular action of RAP as an mTOR inhibitor – specifically its action with the mTORC1 and mTORC2 complexes. According to our studies, our modified RAP molecules outperformed RAP in lower concentrations for inhibiting fibroblast proliferation (mTORC2-mediated) while “dimeric” RAP underperformed in inhibiting migration (mTORC1-mediated). It may be possible that the steric hinder-ance associated with having an additional RAP group flanking the PEG may have interrupt RAP’s complexation with mTORC1. With the conjugation of adamantane, a sterically smaller group, RAP’s function was not attenuated.

While this study showed that the properties of small-molecule drugs can be modified by selectively conjugating select groups, the conjugation yields we reported would pose an obstacle for scaling up the synthesis – around 250 mg of RAP would ultimately yield around 20 mg of purified conjugate. This may have been due to solvent choices, as DMSO:NaHCO_3_ for EDC/NHS conjugation could be further optimization. In addition, as RAP and RAP-Ad had only a 1000 Dalton molecule weight difference, complete purification of one species from the other cannot be assured with dialysis and would benefit from more robust purification methods.

RAP was chosen as a model drug because of its highly sensitive structure-function relationship, however, the proposed strategy could be utilized for any small-molecule drug (or even large-molecule drugs) with the application of improving a variety of suboptimal properties, while maintaining other properties. For example, RAP conjugation to PEG1000 showed to double its solubility in PBS while a single RAP conjugated to PEG3000 tripled its solubility, respectively (*Supp. Fig. 3*). Any carboxylic acid or sterically unhindered alcohol group can be modified in a similar fashion as outlined in this paper. While EDC/NHS coupling could be replaced with higher-yielding ‘click’ chemistries the process of engineering small-molecule bioconjugates based on structure-function analysis is an attractive alternative with advantages beyond random, high-throughput screening. Specifically for RAP, this may allow the creation of an injectable formulation that would not rely on nanoparticles or liposomes for efficient delivery and enhanced solubility in physiological solutions.

In addition, this study presents the opportunity for new combination therapies, as two synergistic drugs could be tethered to one another and simultaneously delivered. For example, RAP and dasatinib combination therapies are becoming common for metastatic breast cancers and could, theoretically, be tethered together to increase colocalization and solubility [30]. Larger payloads, such as proteins or antibodies, could also theoretically be conjugated to one or more “tethers” to allow them to complex with pCD systems, extending their delivery timeframe.

## Conclusion

Our proposed synthesis successfully tethered RAP to either an adamantane group or an additional RAP molecule site-specifically which significantly increased its affinity for cyclodextrin-based hydrogels. This translated to increased loading efficiency and prolonged release from the polymer system, all while maintaining its biofunctionality in vitro. The alteration of a smallmolecule drug’s affinity towards pCD hydrogels displays potential for enhancing other suboptimal drug properties like solubility, delivering larger payloads in affinity-based polymer systems, and extending the window of delivery of small-molecule drugs for long-term therapies.

## ASSOCIATED CONTENT

### Supporting Information

Supplementary Figures (file type, PDF)

## AUTHOR INFORMATION

### Author Contributions

H.A.R. devised the project. A.B.D. developed the technical procedures and performed the experiments under the supervision of H.A.R. Under the supervision of H.A.R, A.B.D wrote the manuscript; all authors read or edited the manuscript.

### Funding Sources

This work was supported by the National Institute of Health (R01 GM121477).

## ACKNOWLEDGMENT

Special thanks to Dr. Nathan Rohner for assisting in cell culture studies and Dr. James Faulk (Department of Chemistry, CWRU) for Department of Chemistry equipment training and usage.

## ABBREVIATIONS

RAP: Rapamycin
pCD: polymerized cyclodextrin
RAP-Ad: RAP-PEG1000-Adamantane
“Dimeric” RAP: RAP-PEG3000-RAP
β-CD: beta cyclodextrin
NMR: nuclear magnetic resonance
FTIR: Fourier Transform Infrared Spectroscopy
SPR: surface plasmon resonance.

## Notes

### Competing Interest Statement

The authors have declared no competing interest.

### Summary of Updates

Enhanced image quality on a handful of figures and minor revisions.

