## Supplemental Figures for "Engineering Selective Molecular Tethers to Enhance Suboptimal Drug Properties" (from another version of this preprint: version 1, 2020-02-02)

Supplemental Materials

**Supp. Fig. 1**

C13 NMR taken to confirm Adamantane conjugation (DMSO-$D_6$).

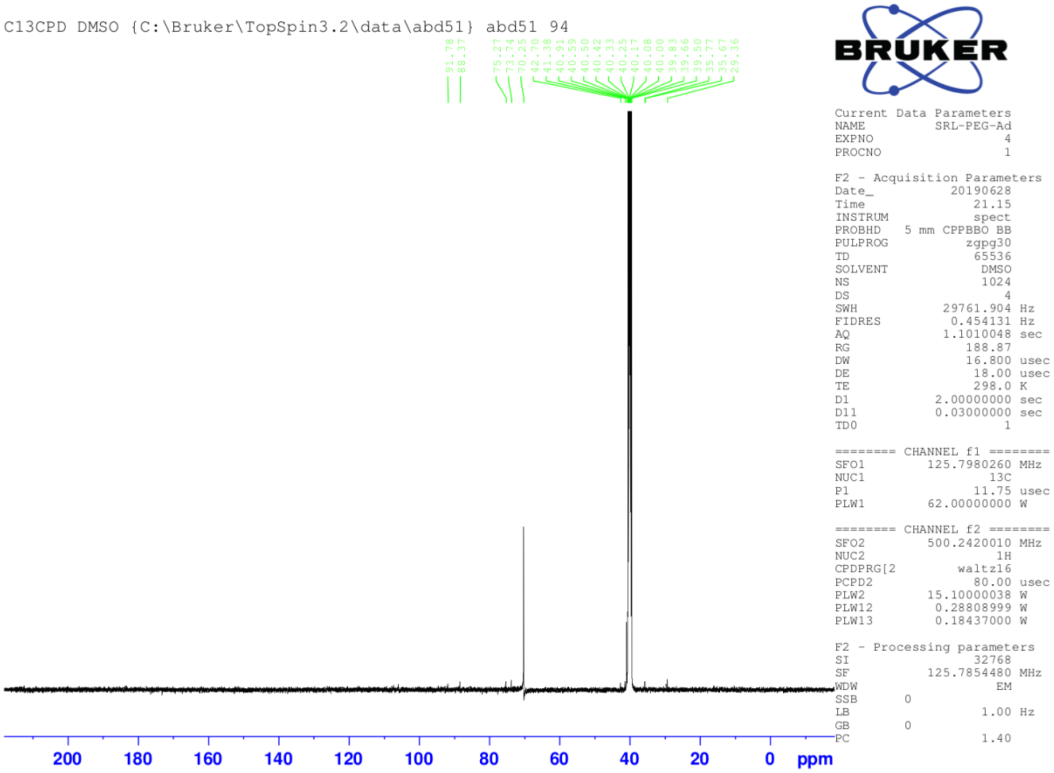


**Supp. Fig. 2**

Non-polar phase HPLC-UV Vis of EDC/NHS Conjugation of Additional RAP. 80:20 v/v ethanol:water eluent. UV-Vis read at 278 nm.

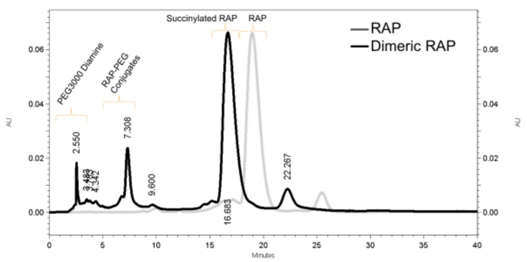


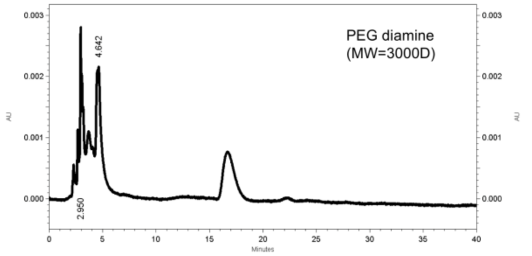


| HPLC UV-Vis Spectrum | Conditions | Product Ratio (%) |
|---|---|---|
| 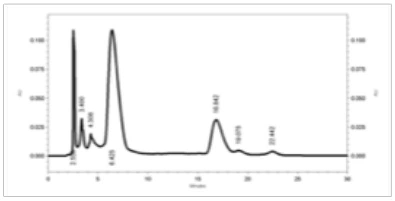 | • Excess EDC/NHS (19/4 mg/ml)<br>• 55% v/v DMSO in $NaHCO_3$ buffer, pH 8.3 | -"Dimeric" RAP / RAP-PEG3000 (39%) *single peak<br>-PEG3000 (51%)<br>-RAP (10%) |
| 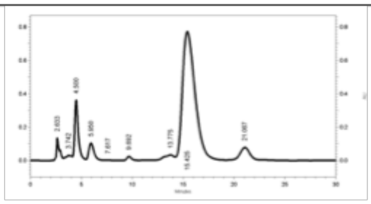 | • 40% v/v DMSO in $NaHCO_3$ buffer, pH 8.3 | -"Dimeric" RAP (2%)<br>-RAP-PEG3000 (8%)<br>-PEG3000 (32%)<br>-RAP (58%) |
| 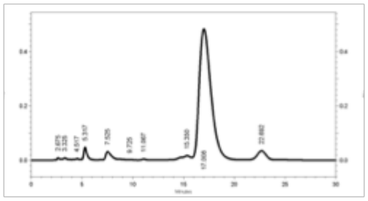 | • 40% v/v DMSO in MES buffer, pH 6.8 | -"Dimeric" RAP (3.2%)<br>-RAP-PEG3000 (2.5%)<br>-PEG3000 (3%)<br>-RAP (80%) |
| 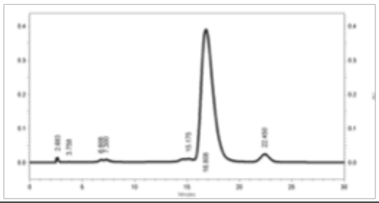 | • Excess EDC/NHS (19/4 mg/ml)<br>• 55% v/v DMSO in MES buffer, pH 6.8 | -"Dimeric" RAP (1.6%)<br>-RAP-PEG3000 (1.7%)<br>-PEG3000 (3.1%)<br>-RAP (86%) |

**Supp. Fig. 3**: Estimated solubility limit of RAP and related compounds in PBS, 1% tween 80. RAP's solubility approached literature value of about 5 uM. Solubility of compound increased with size of PEG conjugated. n=3, 2 batches tested

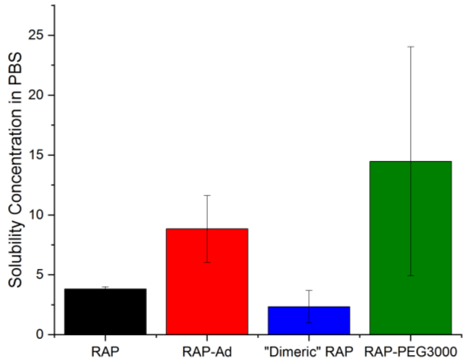
